# Dynamic effects of fructose and *Momordica charantia* supplementation on pulmonary hypertension in broiler chickens

**DOI:** 10.64898/2026.03.05.709939

**Authors:** L.M. Vargas-Villamil, A.C. García-Medina, J.B. Bautista-Ortega, S. Medina-Peralta, L.O. Tedeschi, F. Izquierdo-Reyes, J.M. Zaldívar-Crúz

**Author notes:** For correspondence: Luis Manuel Vargas-Villamil Colegio de Postgraduados, Campus Tabasco, Periférico Carlos A. Molina Km 3.5, Apartado Postal 24, C.P. 86500, Cárdenas, Tabasco, México.

## Abstract

This study evaluated the effects of short-term, low-dose supplementation with high-fructose corn syrup (HFS) and Momordica charantia (Mo) on pulmonary arterial hypertension (PAH) and system dynamics in hypertensive broiler chickens. Thirty male Cobb broilers were divided into three groups (n=10): HFS, Mo, and control. PAH was surgically induced at day 17-19, and supplementation was provided from days 20 to 37. Measurements included body weight, feed intake, heart indices, and tissue composition. A biologically informed dynamic mechanistic model (*Yaantal fe* Bro A1) was used to represent energy utilization and partitioning through the simultaneous calibration of multiple observed trajectories. Correlations between observed and simulated variables indicated generally strong agreement in temporal variation (0.90 < r < 1.0) although these comparisons were based on calibration data. HFS was associated with greater feed intake and weight gain, whereas Mo showed patterns consistent with improved feed efficiency and selective tissue growth. Both supplemented groups showed reductions in the ventricular index used as an indirect indicator of PAH-related cardiac changes, with a greater reduction observed in the HFS group. These exploratory findings illustrate the potential of integrated system dynamics modeling to improve the interpretation of nutritional interventions and inform future experimental designs, potentially supporting the Reduction principle of the 3Rs.

## 1. Introduction

Obesity is a multifactorial metabolic disorder characterized by alterations in adipose tissue metabolism, endocrine dysfunction, and systemic inflammation, often leading to cardiovascular complications such as systemic and pulmonary hypertension. Among the dietary components linked to the increasing incidence of metabolic syndrome, high-fructose corn syrup (HFS) has drawn significant attention due to its pervasive use in processed foods and beverages and its association with insulin resistance, dyslipidemia, and increased adiposity (Bray *et al*., 2004; Stanhope, 2012). Excess fructose intake has been shown to disrupt lipid metabolism, elevate blood pressure, and alter hormonal signals involved in appetite regulation, potentially contributing to the development of both obesity and hypertension (Hwang *et al*., 1987; Johnson *et al*., 2007; Teff *et al*., 2004).

In contrast, *Momordica charantia* (bitter melon), a plant widely used in traditional medicine, has shown promising anti-hyperglycemic, hypolipidemic, and antioxidant properties (Alam *et al*., 2015). Various studies in rodent and human models suggest that *M. charantia* can modulate glucose metabolism, reduce fat accumulation, and improve lipid profiles, supporting its potential as a nutraceutical intervention for obesity-related disorders (Joseph & Jini, 2013). However, comparative studies evaluating the differential metabolic and physiological effects of fructose and *M. charantia* supplementation in controlled experimental settings remain limited.

Pulmonary arterial hypertension (PAH) is a progressive disease that can be studied in broiler chickens because of their susceptibility to pulmonary vascular remodeling and right ventricular hypertrophy under stress conditions (Bautista-Ortega & Ruiz-Feria, 2012). The broiler chicken provides a useful experimental model for investigating pulmonary hypertension and associated cardiovascular alterations, although its findings must be interpreted in light of the differences between animal models and human disease (Colvin & Yeager, 2014). Moreover, its accelerated growth and responsiveness to dietary modulation provide a relevant system for investigating obesity-related metabolic dysfunctions.

The complexity of the metabolic alterations induced by dietary interventions and the need to understand dynamic physiological responses over time call for integrative analytical tools. Dynamic mechanistic modeling, based on differential equations, provides a framework for representing biological processes and examining their responses under different conditions (Tedeschi *et al.,* 2005). The integration of biological knowledge, experimental observations, inverse problems, and multiple-curve fitting can further constrain model behavior and contribute to the understanding of complex animal systems (Vargas-Villamil and Tedeschi, 2014; Vargas-Villamil *et al*., 2020). Here, we propose an integrative approach combining experimentation and modeling to evaluate the physiological and systemic effects of dietary supplementation with HFS and *M. charantia*.

This study employed the ***Yaantal fe* Bro A1** model, a dynamic system developed to represent energy and mass flow across physiological compartments in broiler chickens. This model integrates biologically supported relationships with the simultaneous calibration of multiple observed trajectories to examine tissue-specific growth, energy allocation, and associated cardiovascular responses under dietary and pathological conditions. By integrating experimental observations with mechanistic modeling, this approach seeks to improve the understanding of complex nutritional interactions and the biological processes underlying the observed responses.

### Hypothesis

We hypothesized that low-dose, short-term supplementation with HFS or *Momordica charantia* would produce distinct temporal patterns of feed intake, tissue growth, energy partitioning, and cardiovascular indices, and that the integrated *Yaantal fe* Bro A1 model could describe these physiological responses within a common dynamic framework based on mass and energy balances.

### Objectives

1. To assess the effects of HFS and *Momordica charantia* supplementation on growth performance, feed efficiency, and body composition in hypertensive broiler chickens.
2. To evaluate the impact of these supplements on cardiovascular indices with pulmonary arterial hypertension.
3. To represent energy and mass flows using the *Yaantal fe* Bro A1 dynamic mechanistic model, calibrated simultaneously against multiple observed trajectories, and evaluate the agreement between observed and simulated responses.

## 2. Materials and methods

### 2.1. Study Design

This experimental-modeling study investigated the effects of obesity-related supplementation on hypertension and system dynamics in hypertensive broiler chickens.

#### 2.1.1. Experimental Units and Groups

Thirty male Cobb broiler chicks were randomly allocated to three treatment groups of ten chickens each: a) HFS-supplemented (Fru), *Momordica charantia*-supplemented (Mom) and nonsupplemented control (Tes). Chickens were initially housed together and subsequently allocated to three separate pens, one per treatment group. Supplements were administered individually to the chickens in capsules. Individual chickens constituted biological observational units, while treatment and pen effects could not be separated because each treatment was represented by a single pen. The serial-sacrifice design provided complementary observations at successive sampling times for the reconstruction of treatment-specific dynamic trajectories. This design provided complementary temporal information for mechanistic modeling but did not provide biological replication within each destructive sampling time.

#### 2.1.2. Procedures

Chickens were raised from 1 to 50 days of age. Pulmonary arterial hypertension (PAH) was surgically induced at 17-19 days of age. Supplementation was provided from days 20 to 37. The study was divided into three periods: nonsupplementation (days 1-19), supplementation (days 20-37), and postsupplementation (days 38-50). Supplementation was administered orally once daily from days 20 to 37. *Momordica charantia* leaves were washed, separated, dried, and finely ground. The resulting leaf powder was chemically characterized for dry matter, crude protein, ether extract, neutral detergent fiber, and total polyphenols before encapsulation. Commercial high-fructose corn syrup (HFS) was manually encapsulated, with approximately 0.7 mL per capsule.

The supplementation doses were progressively adjusted according to three body-weight stages. From days 20 to 23 (approximately 450 g body weight), each chicken received 14.4 mg/day of *M. charantia* leaf powder or 0.831 g/day of HFS, according to treatment. From days 24 to 33 (approximately 800 g body weight), the corresponding doses were increased to 28.8 mg/day and 1.669 g/day. Finally, from days 34 to 37 (approximately 1,050–1,300 g body weight), the doses were increased to 57.6 mg/day and 2.484 g/day, respectively. The HFS doses corresponded to one, two, and three capsules per day across these successive stages.

The dosage levels were selected as an exploratory supplementation regimen, considering available information on potential adverse effects and the growth of the chickens. The study was not designed to determine optimal doses or establish dose–response relationships.

#### 2.1.3. Data Analysis

The primary outcome measures were a) body weight and body fraction weights, b) voluntary feed intake, c) excrement output, d) heart-related measurements and e) chemical composition of body fractions. These observations provided information for the descriptive analysis of treatment-specific responses and the dynamic mechanistic modeling described in Section 2.5.

#### 2.1.4. Blinding

Chickens were identified by randomly assigned codes, and researchers conducting measurements and initial data analyses were unaware of the corresponding pen and treatment assignments. The correspondence between animal codes and treatments was disclosed only after the initial statistical analyses had been completed. The sacrifice order was randomized. Animals were randomly assigned to treatment groups using a random number generator.

### 2.2. Research location, housing, and diet

The study was conducted at the poultry facilities of Colegio de Postgraduados, Campus Campeche, México (19°29’54”N 90°32’46”W). Thirty male Cobb broiler chicks were housed from days 1 to 50. The chicks were initially placed in a communal pen for 15 days and then relocated to three pens (1.88 m × 1.30 m × 3.24 m) with wood shaving litter. According to official governmental meteorological records, ambient temperatures during the experimental period ranged from 25 to 28°C. Individual cages (40 cm diameter, one meter high) were used for 12-hour periods to collect individual feed intake and excrement data.

Chickens were fed commercial starter feed (crude protein [CP]= 21.0%; metabolizable energy [ME]= 13,388 kJ/kg) from Days 1 to 29 and finisher feed (CP=19.0%, ME=12,552 kJ/kg) from Day 30 until sacrifice. Water was provided *ad libitum*. Supplements were provided in clear plastic capsules from Days 20 to 37, with doses adjusted according to chicken weights.

### 2.3. PAH induction

PAH was surgically induced in all chickens at 17–19 days of age. The procedure involved occlusion of the external pulmonary artery, as described by Bautista-Ortega & Ruiz-Feria (2012). Chickens were anesthetized via a combination of intramuscular ketamine (HCl, 100 mg mL-1) and xylazine (100 mg mL-1) at a 1:1 ratio, with a dose of 0.01 ml of mixture/100 g body weight. A vascular clip (silver wire, 0.50 mm diameter) was placed on the external pulmonary artery. Postsurgery, the chickens were monitored for two hours under a heat lamp before being returned to their pens. All procedures were performed in accordance with the Guide for the Care and Use of Experimental Animals approved by the General Academic Council of the Colegio de Postgraduados, México.

### 2.4. Body weight, calculated real weight, chemical composition, voluntary feed intake, excrement, body biomass dynamics and thermodynamic flow

Ten chickens per treatment were individually sampled through serial sacrifice, with one chicken sacrificed on each treatment-specific sampling day. Sampling was conducted on days 23, 25, 28, 31, 39, 41, 42, 46, 47, and 48 for Tes; days 21, 22, 24, 29, 32, 35, 38, 39, 44, and 50 for Mom; and days 27, 30, 33, 34, 35, 36, 37, 43, 45, and 49 for Fru. The following measurements were obtained: live body weight (Liv) and the weights of seven body fractions: fat (Fat), breast (Bre), thigh (Thi), viscera (Vis), intestine (Int), bone (Bon), and remainder (Rem). The remainder was defined as the residual body fraction after separation of the other six fractions. Calculated real weight (CRW) was obtained by summing the weights of the seven dissected fractions and was distinguished from directly measured live weight. Individual voluntary feed intake (VFI) and excrement measurements were obtained using the collection procedure described in Section 2.2. Chemical analyses included dry matter, crude protein (CP), ether extract (EE), ash, and total polyphenols.

### 2.5. Model and type of data

A thermodynamic balance-of-mass model, *Yaantal fe* (Tinal-Ortiz et al., 2019), was modified and imported into Stella VI (Doerr, 1996) and Berkeley Madonna software 8.01 (Macey *et al*., 2010). A modified model (*Yaantal fe* Bro A1) was built to evaluate holistic system dynamics, including fraction weight kinetics, and the effects of obesity-related supplementation on the intake, distribution, and deposition of energy in the body system (Figure 1, Supplementary Table_S1_Model_equations.pdf). System dynamics simulations and simultaneous multiple-trajectory parameter fitting were run in Berkeley Madonna software 8.01 (Macey *et al*., 2010).

**Figure 1.**
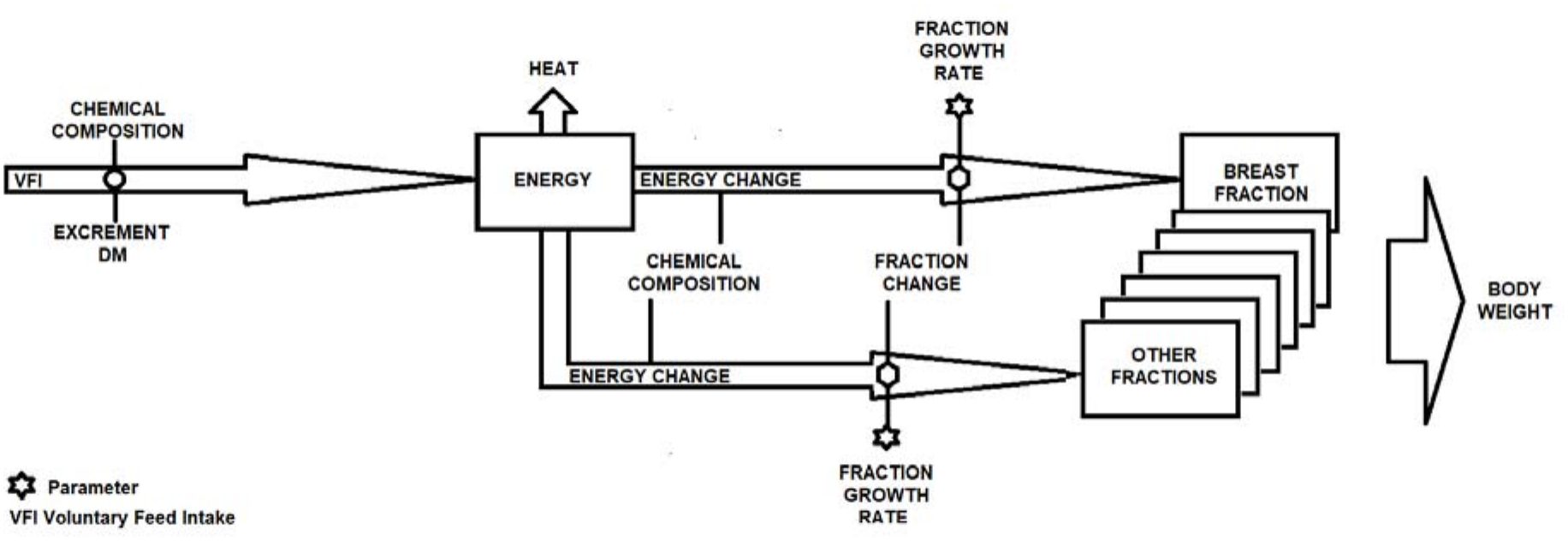
Simplified scheme of the *Yaantal fe* Bro A1 model. Diagram representing the thermodynamic and compartmental relationships used to describe system dynamics and the effect of obesity-related supplementation on hypertensive broiler chickens. The structure reflects intake, deposition, energy flow and biomass distribution pathways. Data modeled and diagram developed by the authors. The definitions of model variables, parameters, and compartment abbreviations are provided in “Supplementary “Table S1 Model equations LV 1011 2026.pdf”.

#### 2.5.1 Dynamic System Modeling (Yaantal fe Bro A1)

The *Yaantal fe* Bro A1 model is a compartmental dynamic system built with differential equations to simulate energy intake, mass allocation, and physiological responses in hypertensive broiler chickens. The system includes first-order differential equations that describe the rate of change in body mass components such as adipose tissue, muscle, viscera, and intestinal mass, as well as cardiac indices.

The model structure incorporates:

- **Energy input (E_1_)** as a function of feed intake.
- **Conversion efficiency (**ε_n_) parameters for each tissue compartment.
- **Growth coefficients (k_i_)** representing tissue expansion over time.
- **Cardiac load functions** modulated by body composition and supplementation effects. The equations follow the general form:

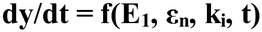

where each compartment’s mass change is driven by intake, efficiency, and regulatory interactions.

Parameter estimation was conducted by simultaneously fitting multiple trajectories of individual observations (Ind) within each experimental group using the Multiple Fits procedure in Berkeley Madonna 8.01, achieving high correlation between simulated and observed data (0.90 < r < 1.0). The Multiple Fits procedure minimizes a combined deviation across multiple variable–dataset pairs, calculated from the root mean square (RMS) deviations of the individual trajectories and their relative weights, as described in the Berkeley Madonna 8.0 User’s Guide (Macey et al. 2010). A complete list of equations and parameters is provided in Supplementary “Table S1 Model equations LV 1011 2026.pdf”.

Three categories of data were distinguished: individual (Ind), mean (Mea), and simulated (Sim). Individual data (Ind) corresponded to measurements obtained from chickens at their respective treatment-specific sacrifice times and constituted the primary observations used for model calibration. Mean data (Mea) represented treatment-level averages of live weight recorded on days 19, 24, 29, 34, 39, and 44, and of feed intake (Itk) and excrement (Exc) recorded on days 20, 25, 30, 35, 40, and 45. These averages included all birds remaining in each treatment group after previous sacrifices; consequently, sample size decreased progressively throughout the experiment. Earlier averages were therefore based on more observations than later ones. Mean data provided complementary descriptive information for comparison with simulated trajectories, while simulated data (Sim) were generated through simultaneous fitting of multiple individual-data trajectories.

#### 2.5.2 Statistical Considerations for Model-Based Design

Due to the nature of the experimental-modeling design, each sampling time represents a single animal per group (n = 1), and the system operates under deterministic principles using a mechanistic dynamic model (*Yaantal fe* Bro A1). Therefore, measures of dispersion (e.g., standard deviation, standard error) or inferential statistics (e.g., p-values, confidence intervals) are not reported per time point, because only one animal was observed per treatment at each sampling time.

To assess whether the *Yaantal fe* Bro A1 model was adequate for evaluating system dynamics, the relationships between observed and simulated variables were investigated using Pearson’s linear correlation analysis. The correlation coefficient (ρ) indicates the strength of association, with values closer to 1 or -1 denoting stronger relationships. SAS version 9.4 was used to generate a correlation matrix. High correlation values indicate agreement in temporal variation between observed and simulated trajectories within the calibration dataset, but do not constitute independent predictive validation. This approach aligns with standard practices in deterministic system modeling (e.g., Berkeley Madonna 8.01), where model validation emphasizes global fit and structural behavior rather than pointwise statistical inference.

### 2.6. Total heart, total ventricular, and right ventricular weights, and API dynamics

The arterial pressure index (API), an indirect indicator of right ventricular changes associated with PAH, was determined in all periods as follows: API = RVW/TVW, where API= arterial pressure index; RVW= right ventricular weight (g), and TVW= total ventricular weight (g). The increases or decreases in the kinetic rates of every heart fraction and API were calculated.

## 3. Results

### 3.1. Feed Consumption, Excrement Dynamics and Chemical Composition

Fru presented the highest voluntary feed intake (VFI) throughout the experiment, particularly during the supplementation and postsupplementation periods (Figure 2). Mom and Tes had similar, lower VFIs. By the end of the experiment, the feed conversion ratios for Fru, Mom, and Tes were 2.31, 1.53, and 1.75 kg of feed/kg animal, respectively.

**Figure 2.**
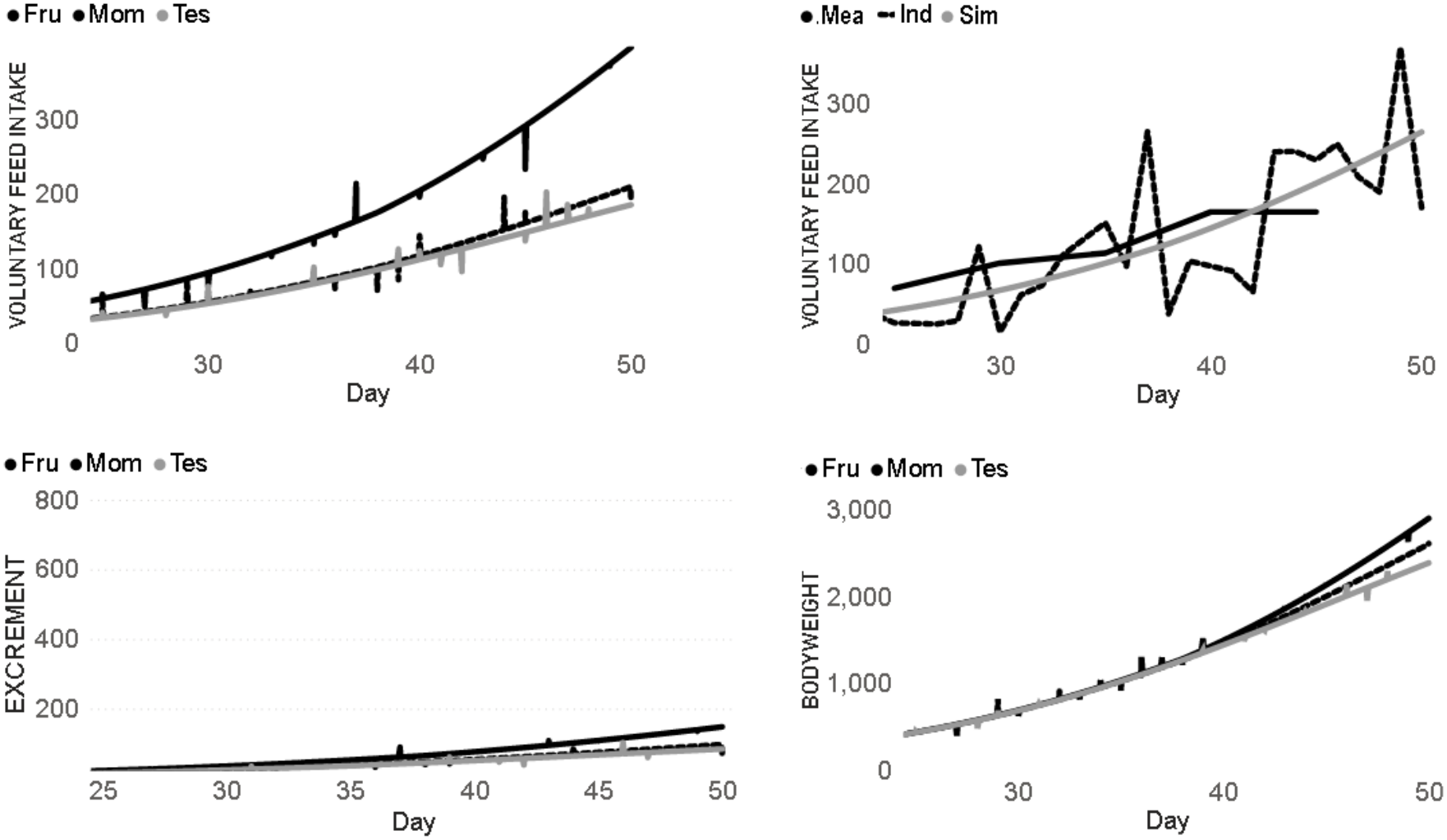
Voluntary feed intake (VFI), excrement (g DM) and body weight (g) dynamics in broiler chickens. Observed and simulated values for VFI (g DM/d), excrement (g DM/d), and body weight (g) in chickens receiving high-fructose corn syrup (Fru group), *Momordica charantia* (Mom group), or no supplementation (Tes group). Data are represented as means (Mea), individual observations (Ind), and simulations (Sim). Data generated by authors. Abbreviations: VFI = Voluntary Feed Intake; DM = Dry Matter.

Excrement production was highest in Fru, with an excrement index of 0.057 g DM/g CRW, whereas it was 0.042 and 0.040 g DM/g CRW for Mom and Tes, respectively (Figure 2).

The chemical composition of the body fractions (Figure 3) showed similar patterns across groups, with a general decrease in compound concentration during growth. An exception was observed in the ether extract (EE) contents of Viscera (Vis) and Intestine (Int), which tended to increase. The fat, bone (Bon), and remainder (Rem) fractions maintained relatively constant compositions.

**Figure 3.**
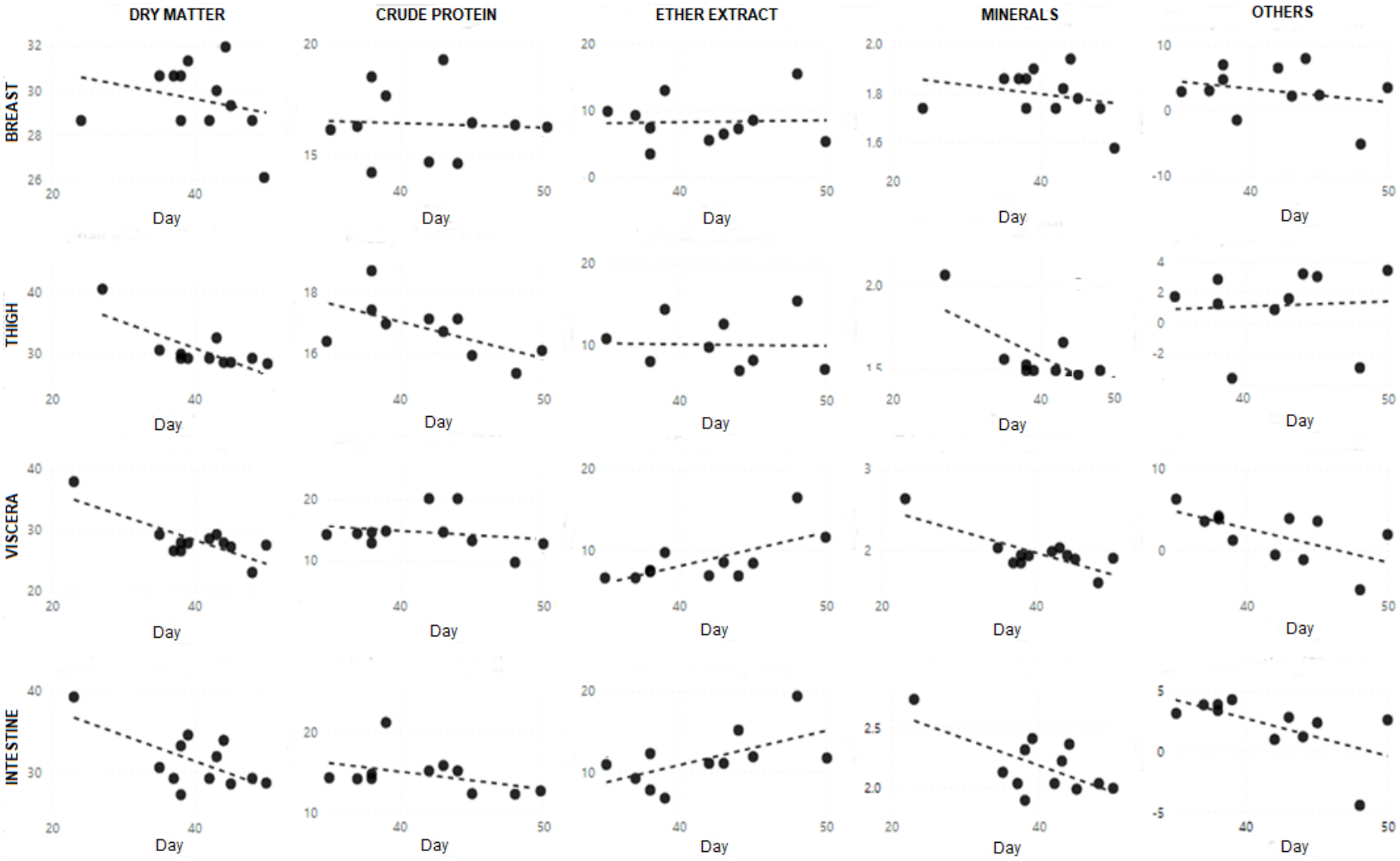
Average chemical composition (%) of body fractions in broiler chickens. Depicts the average ether extract, crude protein, ash, and dry matter content in breast, thigh, viscera, and intestine fractions throughout the experiment. Patterns are shown for the different supplementation groups over time. *Data generated by authors through chemical analysis. Abbreviations: EE = Ether Extract; CP = Crude Protein*.

No outliers were identified by visual inspection within the experimental dataset, and no data points were excluded from the analysis.

### 3.2. Body Weight Dynamics and Fraction Composition

All groups had similar live weights (Liv) and calculated real weights (CRW) that represent the body fraction behaviour (g) during the supplementation period. During the postsupplementation period, Fru presented the highest Liv (Liveweight) and CRW values (Figure 4). The heaviest fractions across all groups were Remainder (Rem), Breast (Bre), and Thigh (Thi), with final weight averages of 680, 578, and 488 g, respectively.

**Figure 4.**
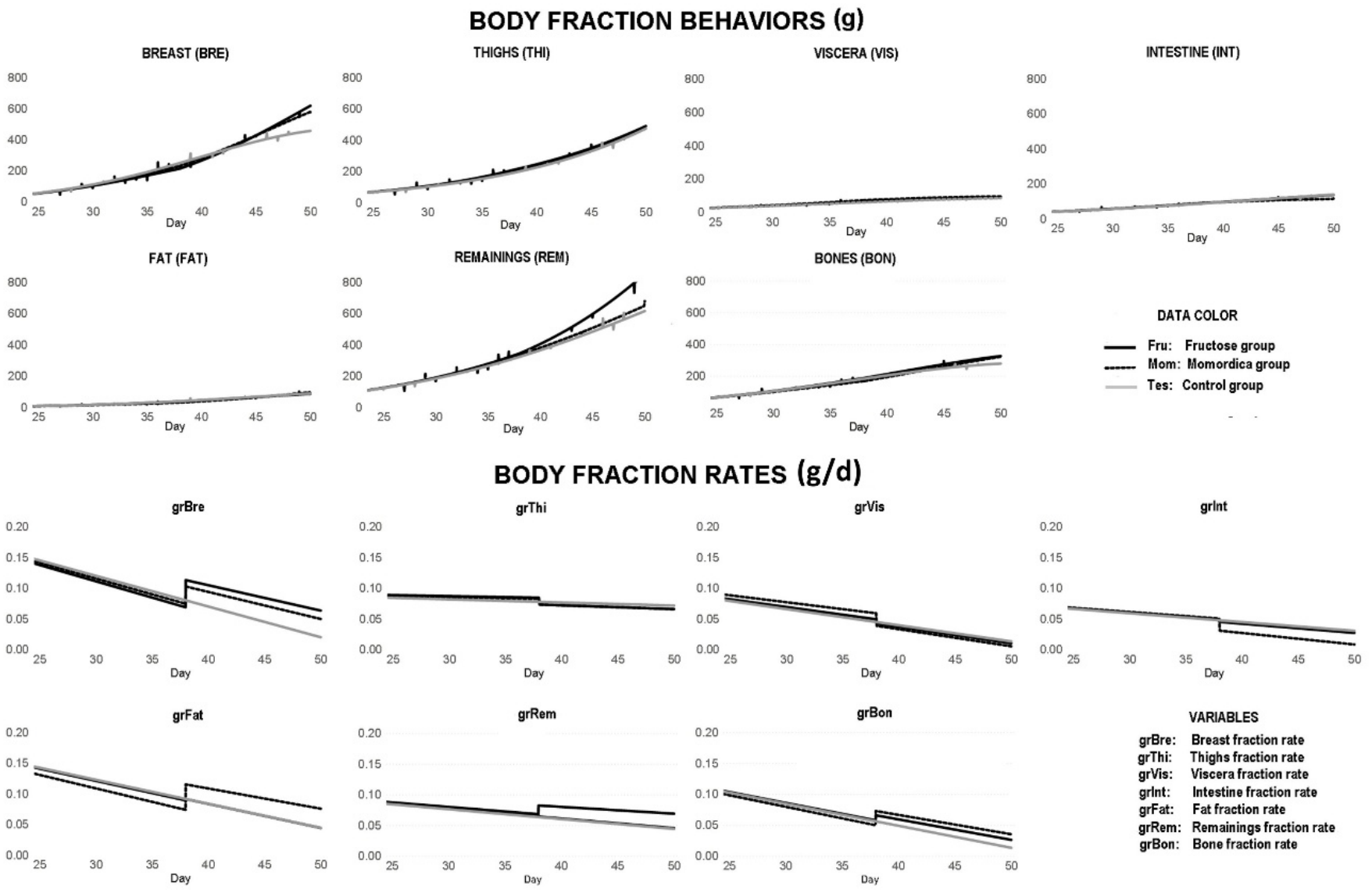
Body fraction behaviours and rates in broiler chickens. Simulated and observed values of body fraction behaviours (g) and rates (g/d) for broilers in each experimental group. Data types include individual (Ind), mean (Mea), and simulated (Sim) observations. *Note. Data generated by authors. Fraction abbreviations: Bre = Breast; Thi = Thigh; Vis = Viscera; Int = Intestine; Rem = Remainder; Fat = Fat; Bon = Bone*.

There was no clear pattern for Int in Fru throughout the experiment. Int in Mom was similar throughout the experiment, but the growth rate of Int (grInt) decreased during the postsupplementation period (Figure 4).

### 3.3. Thermodynamic Flow

Fru presented the highest simulated heat production (Figure 5), which increased continuously until the end of the experiment. Mom and Tes presented lower, similar heat production values (Figure 5). At the end of the supplementation period, heat production for Fru, Mom, and Tes was 692, 63, and 21 J/d, respectively. By the end of the postsupplementation period, these values reached 2,044, 515, and 638 J/d, respectively.

**Figure 5.**
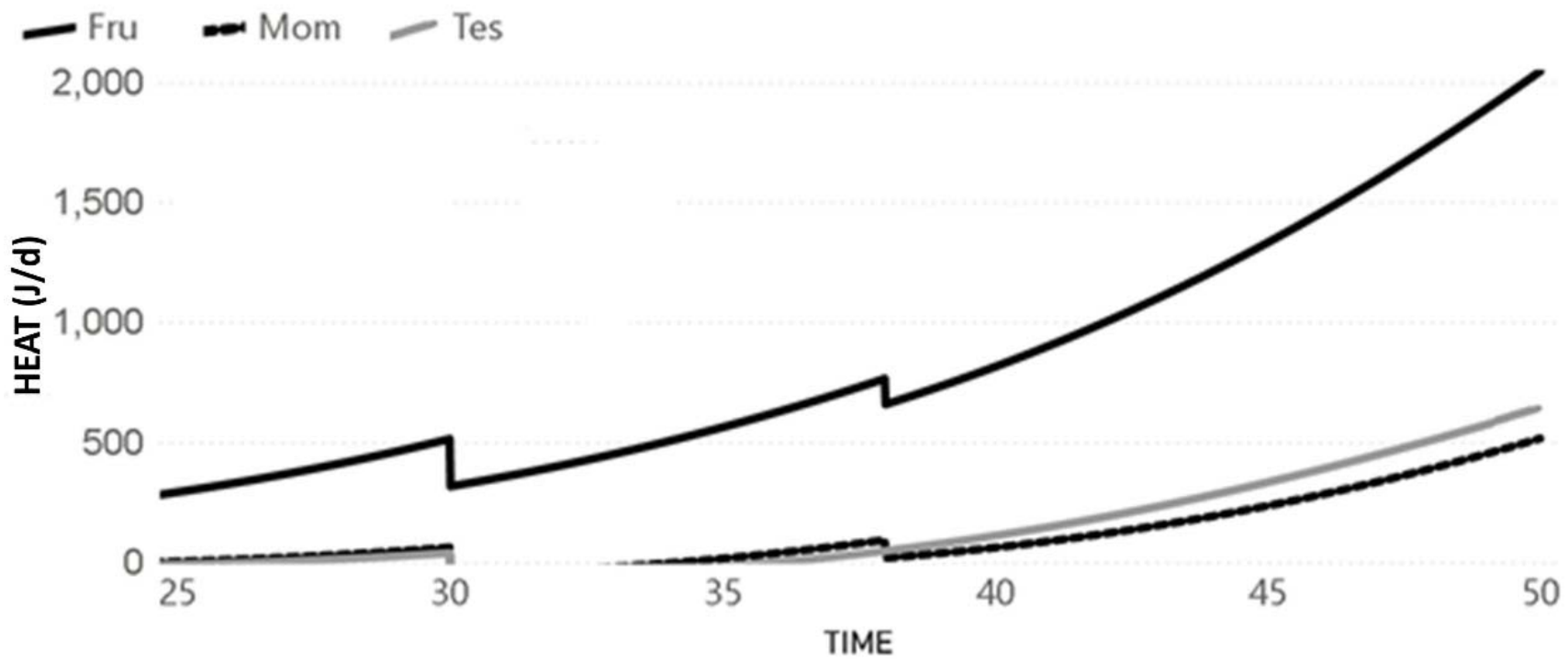
Heat production (J/d) in broiler chickens under different supplementation treatments. Simulated dynamics of heat production derived from energy balance in broilers receiving high-fructose corn syrup (Fru), *Momordica charantia* (Mom), or no supplementation (Tes). *Model simulations performed using Berkeley Madonna software 8.01. Units expressed in Joules per day (J/d)*.

Fru showed the highest feed intake but the lowest energy deposition among the three groups, as indicated by the model-derived energy change (Cha = *Energy change from feed to deposition*) (Figure 6).

**Figure 6.**
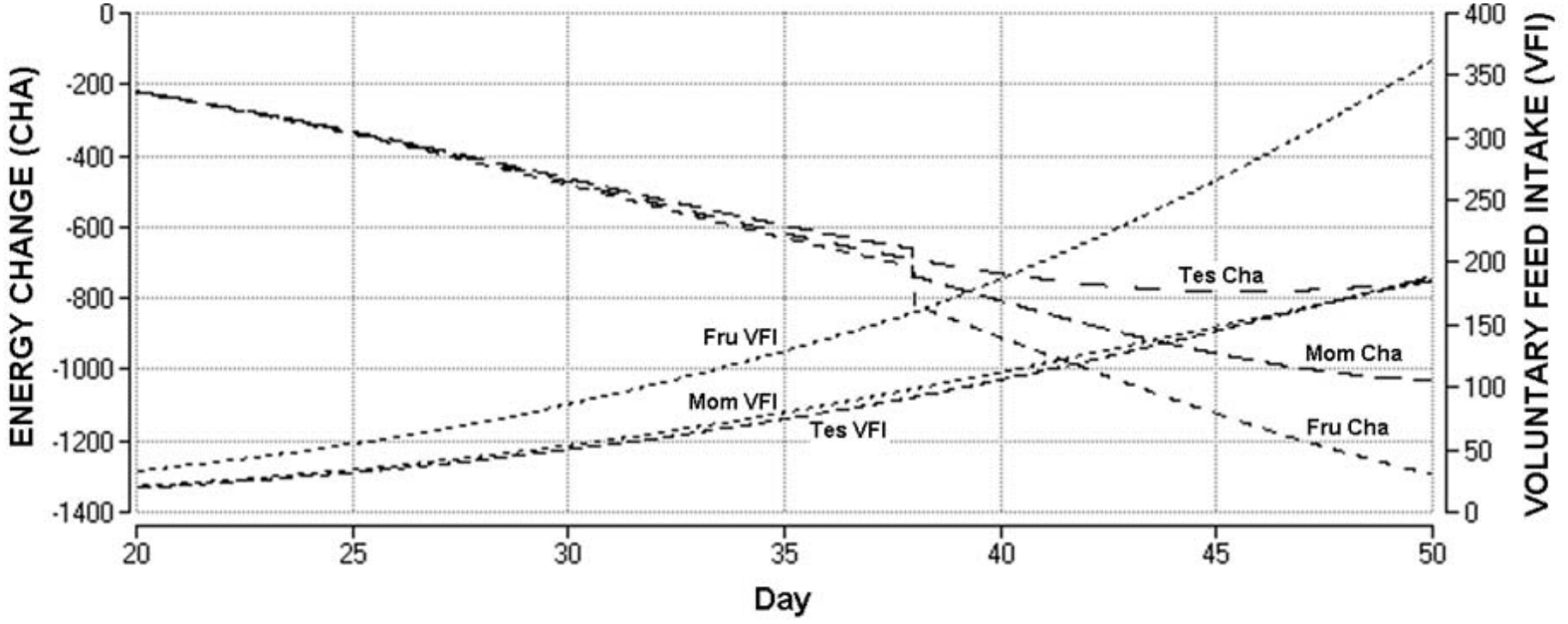
Simulated feed intake and energy change (J/d) across fractions and treatments. Treatments include Fru, Mom, and Tes groups. *Data generated using the Yaantal fe Bro A1 model. Abbreviations: Cha = Energy change (to deposition); VFI = Voluntary Feed Intake*.

### 3.4. Heart Measurements and Arterial Pressure Index

The simulated total heart, total ventricular, and right ventricular weights increased in all groups by the end of the experiment. On Day 50, Fru had the highest total heart (16.38 g) and total ventricular (9.11 g) weights but the lowest right ventricular weight (1.71 g) (Figure 7).

**Figure 7.**
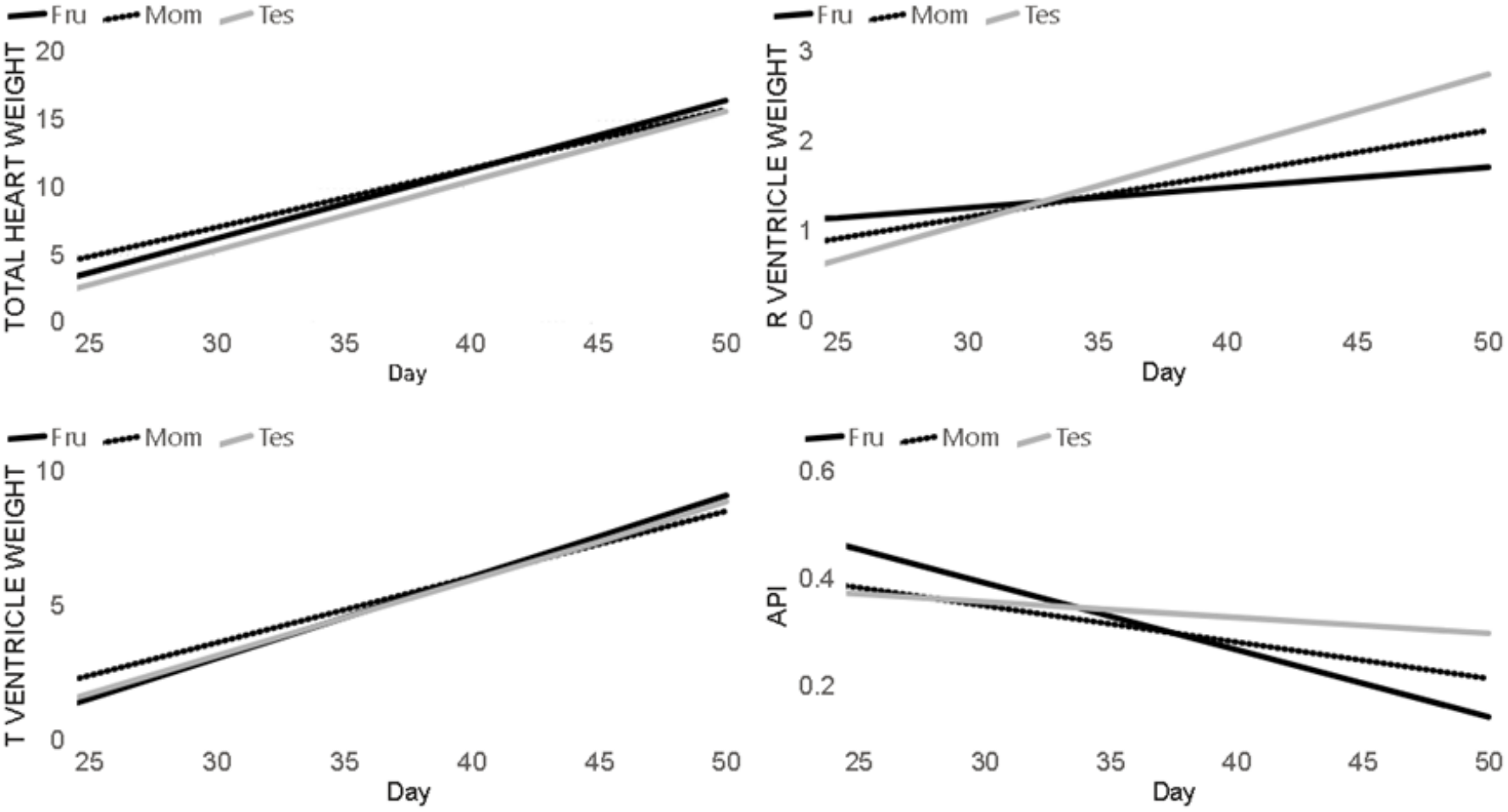
Heart fraction weights and arterial pressure index (API) behavior in broilers. Simulated dynamics of total heart weight, right ventricle weight, total ventricular weight (g), and arterial pressure index across groups and time points. *API = Right ventricle weight / Total ventricular weight. Data generated by authors; linear trends fitted to illustrate temporal patterns*.

The arterial pressure index (API) decreased with age in all groups (Figure 7). On Day 50, the Tes group had the highest API (0.30), followed by Mom (0.21) and Fru (0.14). Relative to their initial API levels, Fru showed the greatest decrease, followed by Mom and Tes (Figure 7).

### 3.5. Model Fit Evaluation

Pearson’s correlation analysis revealed strong associations between the observed and simulated variables for most fractions (0.90 < r < 1), with Fru showing the strongest correlations. The weakest associations were observed for the intake (Itk) and excrement (Exc) variables (0.50 < r < 1). Compared with the other groups, the Mom group generally presented weaker correlations (Figure 8). These correlations describe agreement in temporal variation within the calibration dataset and do not represent independent predictive validation.

**Figure 8.**
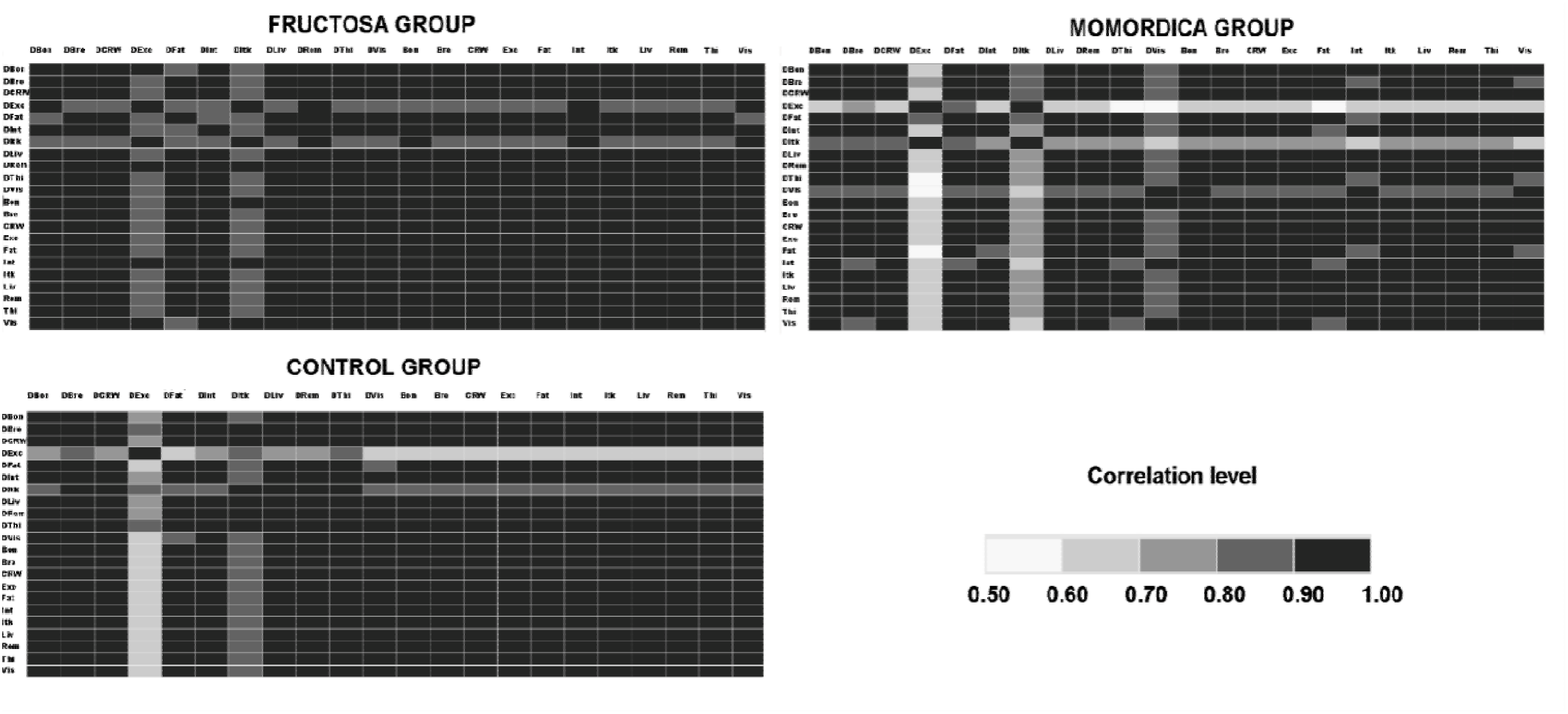
Pearson correlation coefficients between observed and simulated variables. Correlation matrix showing the strength of the relationship between observed (D[Fraction]) and simulated ([Fraction]) values. *Data enerated by authors using SAS software. Abbreviations: Bon = Bone, Bre = Breast, CRW = Sum of fractions, Exc = Excrement, Fat = Fat, Int = Intestine, Itk = Intake, Liv = Liveweight, Rem = Remainder, Thi = Thighs, Vis = Viscera*.

## 4. Discussion

### 4.1. Feed Intake and Appetite Dynamics

The increased VFI throughout the experiment in Fru (Figure 2) could be related to changes in appetite regulation associated with fructose consumption. In mice, Tillman et al. (2014) observed increased caloric intake during high-fructose feeding without evidence of leptin resistance, suggesting that increased intake does not necessarily require impaired leptin signaling. In humans and some animals, the inability of fructose to suppress ghrelin levels, as well as low levels of insulin and leptin in the blood, increases VFI (Teff *et al*., 2004). However, in chickens, ghrelin, which is secreted in the stomach, peripherally and centrally inhibits food intake (Geelissen *et al*., 2006). These differences suggest that mechanisms regulating feed intake may vary among species and in response to different nutrients, limiting the direct extrapolation of findings from other animals to broiler chickens.

There are few data on the effect of Mo on VFI. However, similar VFIs throughout the entire experiment were detected in Mom and Tes (Figure 2). This result is similar to that reported in mice that consumed high-fat diets, although the difference between groups was nonsignificant (Bian *et al*., 2016). Although HFS and Mo both have been reported to reduce plasma glucose and Mo could theoretically stimulate VFI behavior similar to that of HFS, this was not the case. However, whether differences in metabolite signaling, brain responses, or insulin sensitivity contributed to the higher VFI observed in Fru remains to be investigated.

### 4.2. Thermodynamic Partitioning

#### 4.2.1. Energetic intake effect vs. selective effect

Increased weight in Fru could have been due to increased VFI, as was expected, while that in Mom may reflect a more selective pattern of energy allocation toward muscle growth, accompanied by reduced fat accumulation (Figures 2 and 4). In Fru, the weight of almost all the fractions increased compared with those in Tes, suggesting that the weight gain was the result of a nonspecific increase in biomass. The greater body weight observed in Fru was accompanied by increased feed intake and a generalized increase in body fractions. Similar associations between fructose consumption and increased body weight or adiposity have been reported in mice (Rendeiro *et al*., 2015), although the underlying mechanisms may differ between species.

The weight gain in Mom, as suggested by the increase in breast and thigh fractions and reduction in adipose tissue and API, may be consistent with preferential energy allocation toward muscle growth rather than other tissues.

In mice, fructose consumption has been associated with increased body weight and adiposity, together with reduced physical activity, compared with an isocaloric glucose diet (Rendeiro *et al*., 2015). These findings illustrate that dietary carbohydrate composition may influence body mass and energy allocation, although physical activity and energy expenditure were not directly measured in the present study.

In mice fed a high-fat diet, supplementation with M. charantia extracts has been associated with reduced body weight and visceral tissue mass, together with changes in lipid profiles and hepatic antioxidant markers (Wang & Ryu, 2015). These findings provide additional evidence of the metabolic effects of M. charantia, although the specific biochemical mechanisms underlying the responses observed in the present study remain uncertain.

In mice, M. charantia supplementation has been associated with increased whole-body metabolic rate, reduced adiposity, and changes in the expression of genes involved in mitochondrial biogenesis (Lu *et al*., 2013). Although these findings provide a possible metabolic context for the responses observed in the Mom group, the underlying mechanisms were not directly evaluated in the present study.

#### 4.2.2. Feed efficiency and energetic flow

Although Fru had a higher weight, it had a lower feed conversion efficiency and increased ExC index compared with the other groups, suggesting less favorable feed utilization (Figure 2).

However, intestinal permeability, microbiota composition, and the metabolic mechanisms underlying these differences were not directly evaluated in the present study. Mo has been shown to increase fatty acid oxidation and reduce respiratory quotient and serum triglycerides while improving thermogenesis and mitochondrial energy expenditure (Bian *et al*., 2016).

Acute metabolic studies in humans have shown that fructose-containing meals can induce greater postprandial thermogenesis and carbohydrate oxidation than comparable glucose-containing meals (Schwarz *et al*., 1989). However, a review of human trials found no consistent evidence that fructose-containing diets reduce resting energy expenditure over several days, despite the greater thermogenic response observed in acute comparisons (Tappy *et al*., 2013). These findings suggest that the metabolic consequences of fructose intake may depend on the time scale and physiological processes examined; however, energy expenditure was not directly measured in the present study.

### 4.3. Intestinal and Tissue Effects

#### 4.3.1. Intestinal weight and function

Although HFS consumption in rats has been associated with increased intestinal mass due to increased crypt depth and decreased villus height and surface area, this result was not observed in Fru, in which intestinal weight remained constant (Figure 4). One possible explanation is that the duration of exposure was insufficient to produce a detectable change in intestinal weight in broiler chickens. Mo supplementation has been associated with changes in energy metabolism in obese mice (Bian *et al*., 2016). However, whether changes in intestinal absorption contributed to the responses observed in Mom remains unknown, as intestinal absorption was not directly measured in this study.

#### 4.3.2. Visceral, muscle and adipose tissue dynamics

The increased visceral fat weight in Fru may be associated with metabolic responses to fructose consumption, including changes in insulin regulation and sensitivity reported in rodents and humans (Hwang *et al*., 1987; Johnson *et al*., 2007). In contrast, in Mom, the reduction in visceral fat is consistent with the hypolipidemic effects of Mo reported in multiple animal models (Alam *et al*., 2015). Greater growth in the thigh and breast fractions was observed in Mom, suggesting a more selective pattern of tissue growth that may be associated with differences in energy partitioning. In Fru, the generalized increase in body fractions suggests a nonspecific pattern of biomass accumulation, despite the lower total energy deposition across body fractions estimated by the model (Figure 6).

### 4.4 Cardiac and Hypertensive Effects

#### 4.4.1. Heart weight and right ventricular index

In Fru, a higher total heart weight was observed during the postsupplementation period (Day 50), which may be consistent with changes in cardiac growth or ventricular loading conditions, although cardiac hypertrophy and pulmonary arterial pressure were not directly established by the measurements obtained in this study. Nevertheless, the total heart weight in Mom was slightly lower than that in Fru and higher than that in Tes, without a clear pattern indicating cardiac hypertrophy attributable to Mo. In Fru, the increased heart weight was proportional to the body weight, and the arterial pressure index (API) was lower, suggesting differences in the relative contribution of the right ventricle to total ventricular weight. These observations suggest differences in ventricular weight distribution in Fru, although the underlying cardiovascular mechanisms cannot be established from the measurements obtained in this study.

#### 4.4.2. Total and right ventricular weight dynamics

Although Fru presented the highest heart and ventricular weights, the lowest right ventricular weight was observed in this group on day 50, indicating a lower relative contribution of the right ventricle to total ventricular weight. The reduction in right ventricular weight in Fru may be consistent with changes in cardiac loading conditions, although pulmonary arterial pressure was not directly measured. In contrast, Mom presented a moderate increase in right ventricular weight, indicating a different pattern of ventricular weight distribution without establishing changes in cardiac function or pulmonary arterial pressure. These findings may be compatible with the antioxidant, metabolic, and anti-inflammatory effects previously reported for Mo (Alam *et al*., 2015; Bao *et al*., 2013). However, whether these effects contribute to changes in right ventricular remodeling or pulmonary hypertension remains to be established.

#### 4.4.3. Arterial pressure index behavior and treatment impact

API decreased with age in all groups. However, the greatest reduction in API occurred in Fru, followed by Mom and Tes. These findings raise the possibility that the observed reductions in API may be associated with changes in pulmonary arterial pressure or right ventricular loading conditions, potentially involving different mechanisms among groups. In Fru, the observed reduction may be related to changes in vascular volume and metabolic demands. However, these mechanisms remain uncertain, as pulmonary arterial pressure, nitric oxide bioavailability, and endothelial function were not directly assessed in this study. Therefore, the short-term response observed here should not be interpreted as evidence of a direct vasodilatory effect of fructose. Oxidative stress and inflammatory markers have also been investigated in high-fructose-fed rat models treated with M. charantia (Breetha & Krishnamurthy, 2016), although their relevance to pulmonary vascular responses in broiler chickens remains uncertain.

### 4.5. Dynamic Modeling Interpretation

#### 4.5.1. Systemic simulation and fit evaluation

The dynamic system model *Yaantal fe* Bro A1 showed strong correlations between observed and simulated variables, particularly in tissue fractions, thermodynamic outputs, and cardiac indices. The weakest correlations were found in intake and excrement variables, possibly due to daily fluctuations and sampling variability. However, most correlation values were above 0.90, indicating generally strong agreement in temporal variation between observed and simulated trajectories within the calibration dataset, rather than independent predictive validation. The dynamic mechanistic structure of the model allowed for simulation of longitudinal physiological transitions using observations collected at successive time points.

#### 4.5.2. Mechanistic and compartmental interpretation

The model was especially useful in representing differences in tissue-specific growth, energy efficiency, and cardiovascular indices associated with pulmonary hypertension. It also provided model-based estimates of internal variables and processes that were not directly measured, such as energy redistribution, compartment interactions, and time-dependent processes. These estimates depend on the model structure, parameterization, and calibration data and should therefore be interpreted as mechanistic representations rather than independently verified physiological measurements. This level of detail is rarely accessible through experimental data alone and highlights the importance of integrating modeling into experimental animal science. The use of a single-animal-per-time-point design provided complementary observations for reconstructing treatment-specific trajectories and illustrates the potential of mechanistic modeling to inform future experimental designs consistent with the Reduction principle of the 3Rs framework.

#### 4.5.3. Methodological contribution and reproducibility

The transparency of the model is supported by its documented structure and equations, included in Supplementary “Table S1 Model equations LV 1011 2026.pdf”, while computational reproducibility also requires access to the data, parameter values, initial conditions, and calibration procedures. The system of differential equations was used to represent biological processes through simultaneous parameter estimation across multiple observed trajectories and can provide a basis for future scenario exploration. This approach offers future researchers a methodological framework to evaluate metabolic, cardiovascular, or nutritional dynamics in similar models or other species. Additionally, the use of Berkeley Madonna software 8.01provided a flexible and well-established platform for dynamic system modeling in life sciences.

### 4.6. Integrative View and Theoretical Contributions

The present work integrates experimental data and dynamic simulation to analyze the physiological responses associated with HFS and Mo supplementation in a hypertensive animal model. The experimental groups exhibited distinct patterns across multiple physiological variables. The Fru group showed increased feed intake, nonselective tissue growth, greater cardiac mass, and reduced API. Mom showed greater feed efficiency, selective growth of breast and thigh fractions, reduced fat accumulation, and a reduction in API.

Our findings support the idea that systemic modeling can provide insights that go beyond isolated variable analysis, by representing relationships among physiological compartments through energy and mass flows and their associated dynamic trajectories. The study supports the potential application of integrative physiological modeling in experimental designs with ethical limitations and as a basis for future predictive studies requiring independent evaluation. Moreover, it provides a conceptual framework for future comparative studies involving nutraceuticals and metabolic disorders.

### 4.7. Ethical Considerations

This study was conducted under international ethical principles for animal research. The use of a single-animal-per-time-point protocol provided complementary observations for dynamic mechanistic modeling.

This methodological choice illustrates the potential contribution of mechanistic modeling to the Reduction principle of the 3Rs framework, although a reduction in animal numbers relative to alternative experimental designs was not directly demonstrated. Furthermore, the integration of experimental and simulated data allowed additional exploration of physiological trajectories and compartmental relationships without additional animal sacrifice. This strategy is aligned with current trends in ethical animal research that emphasize maximizing scientific output while minimizing animal use.

### 4.8. Final considerations

Obesity is a multifactorial disorder associated with a cluster of simultaneous conditions, increasing the risk of multiple physiological dysfunctions that affect millions of people worldwide. A complete evaluation of supplements associated with metabolic and cardiovascular responses requires diverse methodologies and perspectives. While researchers have focused on the effects of plant supplements, controlled diets, genetics, metabolic factors, clinical factors, and energetics, integrating these processes within dynamic mechanistic models remains an important methodological challenge, including in experimental animal systems.

The integrative dynamic mechanistic *Yaantal fe* Bro A1 model used in this work, which showed generally strong agreement between observed and simulated trajectories within the calibration dataset, helps improve the understanding of the biological system as a whole by representing physiological relationships through energy and mass flows. This approach lays a foundation for advancements in experimental modeling methodologies, as suggested by Carneiro *et al*. (2019) and Carvalho *et al*. (2021) in their studies on residual feed intake and carcass characteristics.

The productive behavior and body fraction trajectories differed between Fru and Mom, with Fru showing a more generalized increase in body fractions and Mom showing more selective changes in tissue growth. During the postsupplementation period, these differences became more apparent in the trajectories of individual body fractions. The high live weight (Liv) in Fru during the postsupplementation period was associated with markedly high VFI and changes in the growth of specific body fractions, although the underlying metabolic mechanisms could not be established from these observations. Previous studies have described relationships between fructose consumption and metabolic regulation (Tillman *et al*., 2014; Stanhope & Havel, 2008).

Unexpectedly, Fru showed a greater reduction in the arterial pressure index (API) than Mom, raising the possibility of differences in pulmonary vascular or right ventricular loading responses, although pulmonary arterial pressure was not directly measured. These findings appear to contrast with those of Johnson *et al*. (2007) in humans and Hwang *et al*. (1987) in rats. However, further long-term evaluations are needed to determine whether the short-term responses observed in this study persist over time and to clarify the mechanisms involved. Previous research has examined the metabolic effects of prolonged dietary fructose exposure in rats (Tuovinen & Bender, 1975).

## 5. Conclusions

Fru showed greater body weight gain but reduced feed efficiency, whereas Mom exhibited a more selective pattern of tissue growth, particularly in breast and thigh fractions, together with improved feed efficiency and lower fat accumulation. These responses may have practical implications for poultry production, although their economic significance requires further evaluation.

This study found that short-term, low-dose supplementation with high-fructose corn syrup and *Momordica charantia* was associated with reductions in the arterial pressure index (API) in broiler chickens with surgically induced pulmonary arterial hypertension, with distinct patterns of physiological responses between groups. Although Fru showed the greatest reduction in API, this finding may reflect a short-term compensatory response rather than a sustained cardiovascular benefit. Because pulmonary arterial pressure was not measured directly, and prolonged fructose exposure may have adverse metabolic effects, longer-term studies are required to clarify the physiological significance of this response.

The experimental-modeling approach using the *Yaantal fe* Bro A1 system provided a dynamic mechanistic representation of energy and mass flows, with generally strong agreement between observed and simulated trajectories within the calibration dataset. By integrating the three treatment responses into a single dynamic model, this approach maximizes the information obtained from the experimental animals and may support the Reduction principle of the 3Rs by informing future experimental designs. However, its potential to reduce animal use requires further evaluation and independent model validation.

These findings contribute to the understanding of nutritional interventions in broiler chickens with experimentally induced pulmonary hypertension and support the broader use of system dynamics in animal science research.

## 6. Data availability statement

The data and supporting materials are openly available in Figshare at https://doi.org/10.6084/m9.figshare.26207393. These materials include the experimental dataset, the Welfare Committee letter, graphical abstract, Supplementary “Table S1 Model equations LV 1011 2026.pdf” containing the model equations, and a summary document. The repository also provides the Stella 10.1 files in which the *Yaantal fe* Bro A1 model was developed, the Berkeley Madonna files used for model calibration, including the model and fitting parameters, and the Power BI Desktop files containing the data values and graphical representations used in the study. These files are publicly accessible and can be downloaded without access restrictions. Their examination and execution require the corresponding software and an appropriate level of mathematical and modeling expertise. The software applications themselves are not distributed as part of this study.

As an additional means of accessing and examining the model, a cloud-based environment using AWS AppStream is available upon request for scientific and research purposes. This environment provides access to Berkeley Madonna 8.01, Stella 10.1, and Power BI Desktop 2026, together with the corresponding project files, allowing researchers to examine the model structure, equations, parameters, calibration, and results without installing these applications locally. Access requires identification of the requesting researcher and a description of the intended scientific use. Because of operating costs and limited computational capacity, access is subject to availability and may involve waiting periods. This supplementary service does not restrict the public availability of the files deposited in Figshare. Researchers interested in accessing the AWS AppStream environment should contact the corresponding author by email for further information and instructions on the access procedure.

## 9. Graphical Abstract or Cover Image Note

The graphical abstract (cover image) included in this submission was not generated by artificial intelligence.

## 10. Institutional review Board (Welfare Committee) statement

The study was conducted in accordance with the institutional guidelines for the care and use of experimental animals and was approved by the Welfare Committee of Colegio de Postgraduados, México (protocol number: COBIAN/006/21; approval date: October 5^th^, 2021).

## 11. Institutional review Board (Graduate Committee) statament

The graduate protocol was reviewed. Recommendations for improvement were issued, and incorporated, and the project complied with the General Regulations of the Colegio de Postgraduados (2015), and the Curriculum Plan of Graduate program, (PROPAT).

## 13. Author contributions

**L. M. Vargas-Villamil**: Conceptualization, Methodology, Software, Formal analysis, Investigation, Supervision, and Writing – review & editing.

**J. Bautista-Ortega**: Conceptualization, Methodology, Investigation, Supervision, Project administration, and Writing – review & editing.

**L. O. Tedeschi**: Methodology and Writing – review & editing.

**F. Izquierdo-Reyes**: Methodology, Software, Validation, Formal analysis, and Writing – review & editing.

**S. Medina-Peralta**: Methodology, Investigation, and Writing – review & editing.

**A. C. García-Medina**: Investigation, Data curation, and Writing – original draft.

**J. M. Zaldivar-Cruz**: Investigation, Supervision, and Writing – review & editing.

All authors have read and approved the final version of the manuscript and agree to be accountable for their respective contributions and for ensuring that questions related to the accuracy or integrity of the work are appropriately investigated and resolved.

